# Pan-cell type continuous chromatin state annotation of all epigenomes from the International Human Epigenome Consortium

**DOI:** 10.1101/2025.02.06.636950

**Authors:** Habib Daneshpajouh, Ismail Moghul, Kay C Wiese, Maxwell W Libbrecht

## Abstract

The International Human Epigenome Consortium has generated thousands of epigenomic datasets that measure various biochemical activities in the genome, including transcription factor binding, histone modification, and DNA accessibility. Currently, the predominant methods for integrating these datasets to annotate regulatory elements are segmentation and genome annotation (SAGA) algorithms. The majority of annotations by these methods are cell type-specific. However, as the number of profiled cell types has grown into the thousands, using thousands of cell type-specific chromatin state annotations proves undesirable for many applications. Here, we present a pan-cell type annotation that summarizes all IHEC epigenomes using the recently-developed method, epigenome-ssm.

## 1 Background

Annotating the regulatory elements in the human genome helps better understand the mechanistic basis of genetic disease. To this end, international projects including ENCODE, Roadmap Epigenomics, BLUEPRINT, CEEHRC, DEEP, GIS, AMED-CREST, and EpiHK have generated thousands of epigenomic datasets such as ChIP-seq, DNase-seq, and ATAC-seq that measure various biochemical activities in the genome, including transcription factor binding, histone modification, and DNA accessibility. More recently, the International Human Epigenome Consortium (IHEC) unified these projects in order to re-process all these epigenomic data sets through a uniform pipeline. The IHEC repository contains histone modification ChIP-seq, gene expression RNA-seq, and DNA methylation WGBS data across hundreds of human biosamples.

Integrative analysis of these data sets is crucial to understanding the regulatory landscape of the human genome and its role in phenotype and disease. Currently, the predominant methods for integrating epigenomic datasets to annotate regulatory elements are segmentation and genome annotation (SAGA) algorithms such as ChromHMM [1] and Segway [2]. SAGA algorithms take as input data from a given cell type, partition the genome, and assign a chromatin state label to each segment indicating the epigenetic activity at that position [3–27]. Consequently, they can identify regulatory elements without requiring prior knowledge of known genomic elements.

As the number of profiled cell types has grown into the thousands, using thousands of cell type-specific chromatin state annotations proves undesirable for many applications. Therefore, recently, researchers have sought a single unified annotation of regulatory elements across all cell types, known as a “pan-cell type”, “universal”, or “stacked” annotation [7, 13, 28, 29]. A pan-cell type annotation summarizes the activity of each genomic position across all profiled cell types and thus is ideal for understanding evolutionary conservation or disease association. However, pan-cell type annotation is challenging because the combinatorial number of patterns of activity may grow exponentially in the number of cell types.

To tackle this challenge, we recently developed epignome-ssm [30], a method that summarizes each genomic position as a vector of continuous chromatin state features, in contrast to the previous approach of assigning a discrete chromatin state label. This continuous approach can summarize complex high-dimensional datasets into a small number of interpretable chromatin state features. Unlike discrete labels, these continuous features preserve the underlying continuous nature of the epigenomic signal tracks, can easily represent varying strengths of a given element, can represent combinatorial elements with multiple types of activities, and are shown to be useful for expressive visualizations because they map complex high-dimensional datasets onto a much smaller number of dimensions.

Here, we present a chromatin state feature map summarizing all IHEC epigenomes. This feature map comprises 33 genome-wide chromatin state feature signal tracks that summarize all data sets. Each feature track captures a regulatory program that drives gene expression in one or more cell types. For example, as described below, one of the feature tracks (referred to as the Prom-Brain feature) captures the regulatory activity of the promoters active in the brain-related cell types. Together, these features provide a compact representation of chromatin states that is conceptually distinct from traditional discrete genome segmentations, and our framework offers a principled way to integrate, interpret, and label them across thousands of signal tracks with mnemonics that jointly reflect activity type and cell type specificity.

We show that these feature maps constitute an intuitive and visualizable summary of epigenomic data, and that they enable accurate identification of mechanisms of disease association. We show that this is advantageous over alternative pan-cell type mapping approaches according to several metrics. Thus, we expect that this pan-cell type chromatin state feature map will provide a crucial resource to the community of investigators using IHEC data, and enable many downstream analyses and insights.

## 2 Results

### 2.1 Pan-cell type chromatin state feature maps across thousands of human epigenomes

We present a continuous chromatin state feature map generated using epigenome-ssm on 9,539 genome-wide signal tracks from six core histone modification assays (H3K4me1, H3K27ac, H3K4me3, H3K36me3, H3K27me3, and H3K9me3) across 1,698 epigenomes (Fig. 1A, Methods). Briefly, epigenome-ssm takes as input 9,539 genome-wide signal tracks and outputs 33 pan-cell type chromatin state feature tracks (Fig. 1B-C, Methods). Each feature track summarizes a subset of the input tracks and captures a regulatory program that drives gene expression in one or more cell types.

**Figure 1:**
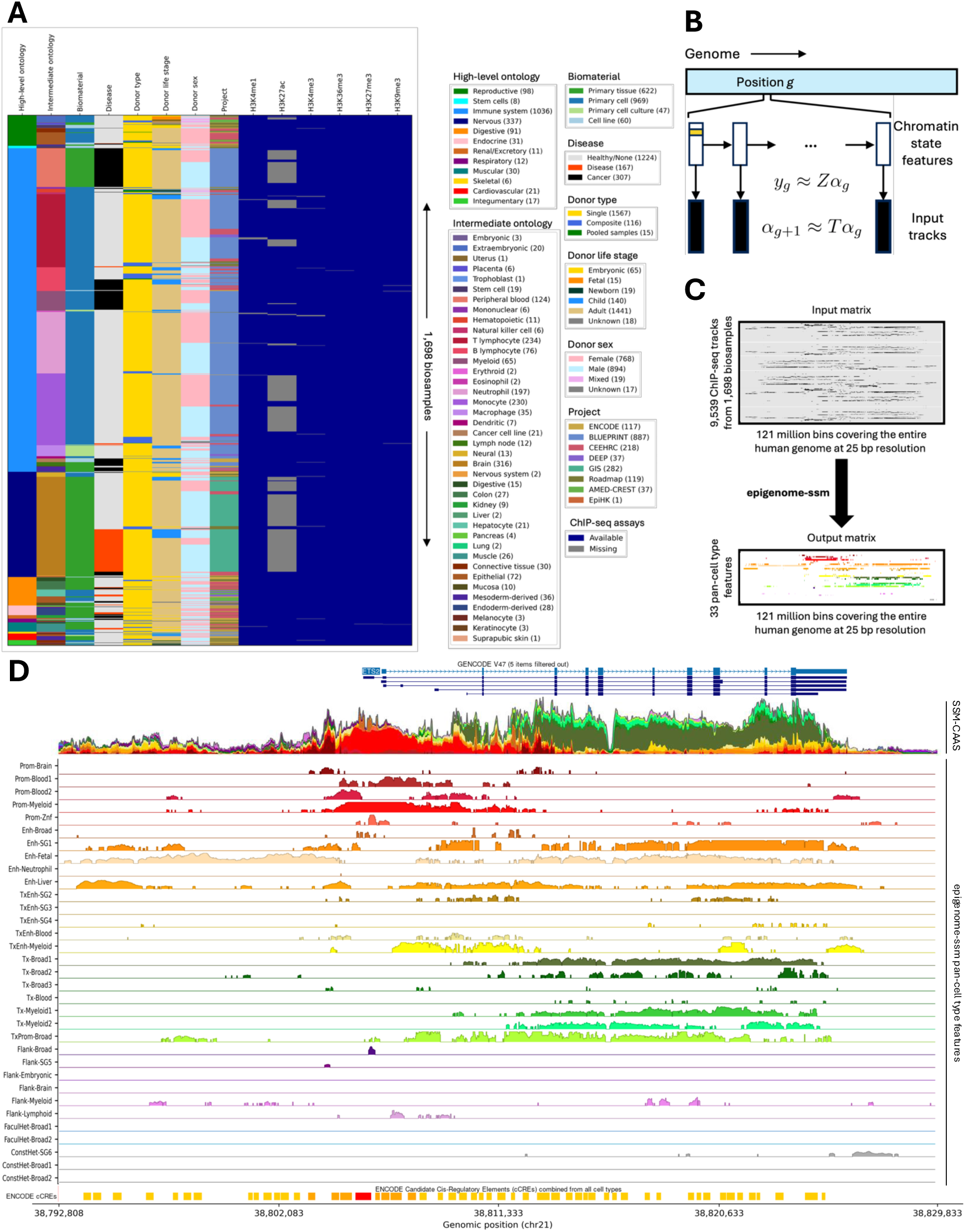
We used epigenome-ssm to generate 33 pan-cell type features across 1,698 IHEC biosamples. **A** The IHEC dataset contains signal tracks from six core histone modification ChIP-seq assays across 1,698 biosamples. **B** epigenome-ssm uses a state space model in which chromatin state features at each position of the genome are modeled by an emission matrix *Z* and a transition matrix *T* (section 5.2 and [30]). **C** A total of 9,539 genome-wide tracks are passed to epigenome-ssm as input. It outputs a vector of *K* chromatin state features at each genomic position that represents the pan-cell type activity at that position. Here, we present a model with 33 chromatin state features. **D** UCSC Genome Browser visualization of pan-cell type features at the region around ETS2 gene (see Fig. 5 for color labels). See Methods section 5.11 for details on how the SSM-CAAS track is generated.

The epigenome-ssm model is an unsupervised state-space model. It assumes that there is an unobserved vector of nonnegative chromatin state features at each genomic position, and that observed signal tracks are generated as a linear function of these chromatin state features. We trained the model using an expectation-maximization algorithm to learn parameters and active set optimization to enforce nonnegativity constraints, iterating until it reached a stable log-likelihood. After training, we normalized the feature values by their genome-wide means to enhance interpretability and standardize values for downstream analyses. This approach is similar to stacked SAGA methods like ChromHMM and Segway, with a few differences, except that existing SAGA methods generate a single discrete label per position, whereas epigenome-ssm generates a vector of 33 continuous values.

As an illustrative example, consider the ETS2 locus (Fig. 1D). This gene is on the forward strand and has multiple transcription start sites (TSSs), and is known to be expressed in several blood cell types, especially the myeloid lineage of the immune cells, such as monocytes and neutrophils (Fig. 1D, GENCODE track). The continuous pan-cell type features provide a more complete picture of the regulatory activities at this locus than both the GENCODE gene annotation and ENCODE candidate cis-regulatory elements (cCREs). epigenome-ssm features show several blood-related promoter features that are enriched around the promoter of ETS2. The Prom-Blood1 feature captures the inner TSSs, Prom-Blood2 captures the most 5’ TSS, and most importantly, Prom-Myeloid is highly enriched around all the TSSs. As shown in the ENCODE cCRE track, this gene contains several intronic enhancers; these are captured by the Enh-SG1, Enh-Fetal, Enh-Liver, and TxEnh-Myeloid, which are enriched for different subsets of those enhancers. Finally, three transcription-related features, Tx-Myeloid1, Tx-Myeloid2, and TxProm-Broad capture the pattern of transcription of this gene across different cell lineages.

We also found that epigenome-ssm features are complementary to the existing pan-cell type annotations such as the ENCODE cCREs (Fig. 1D, bottom track). The ENCODE cCRE annotations contain the position of candidate enhancer and promoter elements. ENCODE cCREs were generated based on the combination of different epigenomic signals in each region. However, such annotation may not provide a complete picture of all the epigenomic activity in a given locus. For example, the two inner TSSs are annotated as enhD (i.e., distal enhancer), while those are likely combinatorial regions exhibiting both enhancer and promoter activities.

One of the design considerations when training unsupervised annotation models is the number of labels (or the number of features in the case of continuous chromatin state models). Here, we initially trained the model with potentially more features than needed (i.e., with 50 features). Then, we sorted all the features by their total emission value, corresponding to each feature’s overall contribution to the model, and used an elbow method to drop 17 features with low total emission (Fig. S1). This approach is analogous to the common practice used in Principal Component Analysis (PCA), where the principal components with the largest eigenvalues explain the most variance in the data. Just as we often discard PCA components with low eigenvalues to retain only the most informative ones, we retained 33 pan-cell type features with high emission values, ensuring that our model focuses on the most meaningful chromatin states while minimizing redundancy.

### 2.2 Pan-cell type features outperform alternatives according to quantitative metrics

We hypothesized that a good annotation should capture the key genomic activities involved in gene regulation, so we assessed the quality of the pan-cell annotations by examining their associations with gene expression, enhancer activity, and evolutionary conservation. Specifically, for each method and each genomic activity, we trained a linear regression model that used the annotations as predictors to estimate the corresponding activity levels.

We compared epigenome-ssm to alternative pan-cell type annotations (Methods). Five methods are compared: DHS-NMF with 16 features [28], Sei with 40 sequence classes [31], Universal ChromHMM with 100 states [29], and epigenome-ssm with both 16 and 33 features. Across all tasks, the epigenome-ssm models consistently achieve a high prediction performance. We found that all the methods are predictive of gene expression. However, epigenome-ssm annotations led to better prediction performance using a smaller number of features than the alternative methods (Fig. 2A-D, supplementary Fig. S2A-D). Notably, even with just 16 features, epigenome-ssm rivals or exceeds the performance of ChromHMM, which uses 100 discrete states, highlighting the efficiency and predictive power of the continuous features.

**Figure 2:**
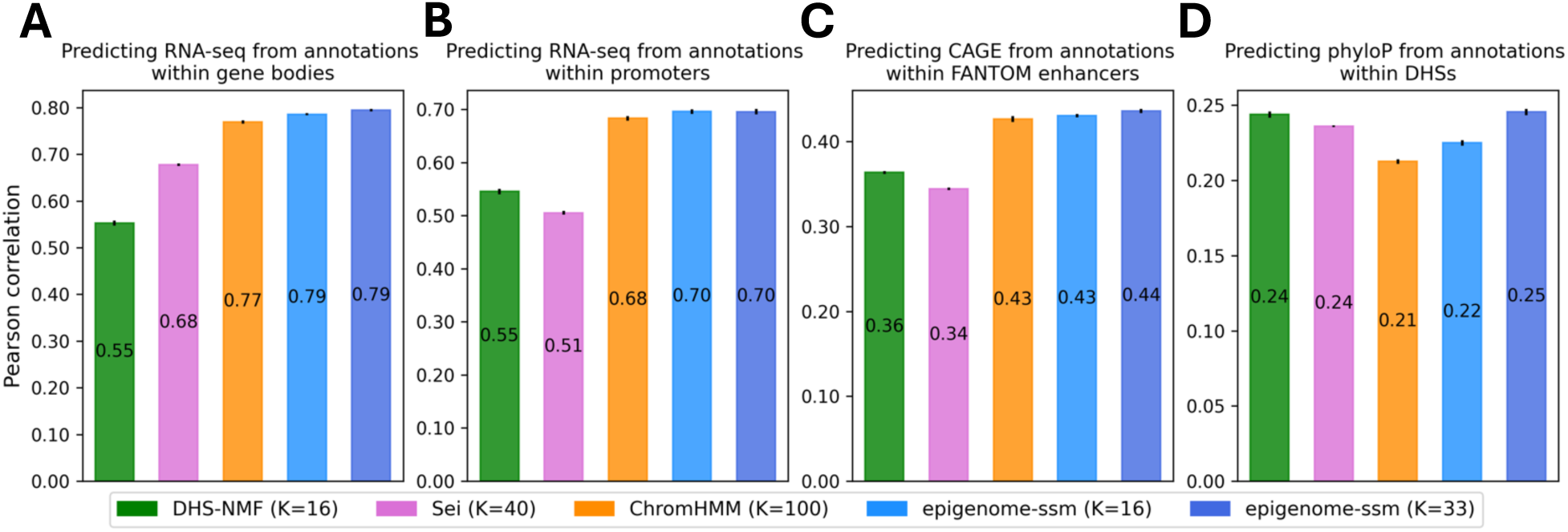
Pan-cell type features are predictive of genomic activities. (A-D) Evaluating pan-cell type annotations according to their prediction power of various genomic phenomena. *K* is the number of components, sequence classes, labels, and features for DHS-NMF, Sei, ChromHMM, and epigenome-ssm, respectively. The vertical solid lines at the top of the bars represent the standard error from a 10-fold cross-validation. **A** Using annotations from within gene bodies to predict RNA-seq TPM values across all epigenomes (see section 5.4). **B** Same as **A** but using annotations within a 2 kb window centered at TSS. **C** Using annotations from within FANTOM5 enhancers to predict CAGE TPM values across all samples (see section 5.5). **D** Using annotations from within DHSs to predict evolutionary conservation, as measured by phyloP (see section 5.6).

We also found that the performance of epigenome-ssm is robust to changes in the number of features (K) (Fig. 2A-D, supplementary Fig. S2A-D). Both the K=16 and K=33 models yield similar correlations, with slight improvements when using more features. This indicates that our pan-cell type features capture the essential aspects of the epigenomic data, providing reliable predictions across different genomic activities with a relatively small set of features.

The better performance of epigenome-ssm highlights the better expressive power of continuous features compared to the discrete chromatin state labels. For example, the intensity of a promoter feature tends to correlate with the expression level of genes regulated by promoters of that type (Fig. S2E). In discrete SAGA, the differences between low- and high-intensity regulatory elements is often represented with separate labels of e.g. “Weak Enhancer” and “Weak Transcription”, which makes analysis challenging [1, 7].

Additionally, we compared epigenome-ssm with CSREP [32], a method for summarizing SAGA annotations across many cell types, and found that the pan-cell type features yield a better prediction performance compared to CSREP (Fig. S3). CSREP summarizes SAGA annotations from a group of related samples into a single annotation that summarizes the group. A collection of CSREP annotations is similar to a pan-cell type annotation in the sense that both reduce thousands of experiments to a manageable size. We compared epigenome-ssm to CSREP summaries of 1,698 IHEC samples down to 56 summary annotations.

### 2.3 Pan-cell type features capture a variety of types of epigenomic activities

We found that epigenome-ssm chromatin state features encompass the known diversity of epigenomic activities. Each chromatin state feature captures both a type of activity and its pattern of activity across cell types. Thus, where a cell type-specific annotation of cell type X might have a mnemonic “Enh” meaning “enhancer active in cell type X”, a pan-cell type model feature has mnemonics of the form “Enh-Blood”, meaning “enhancer active across blood cell types”. Therefore, the mnemonics that we assigned to each pan-cell type feature represent a pair of activity and cell type. We found several main categories of features (Table 1, Fig. 3) by looking at features from different perspectives: (1) Model emissions that associate features with the input tracks with respect to different characteristics of the input samples such as assay, ontology, and donor information; (2) Average feature value at annotated genomic elements such as the transcription start sites, DNase I hypersensitive sites (DHSs), introns and so on; (3) Gene expression; and (4) Gene Ontology (GO).

**Figure 3:**
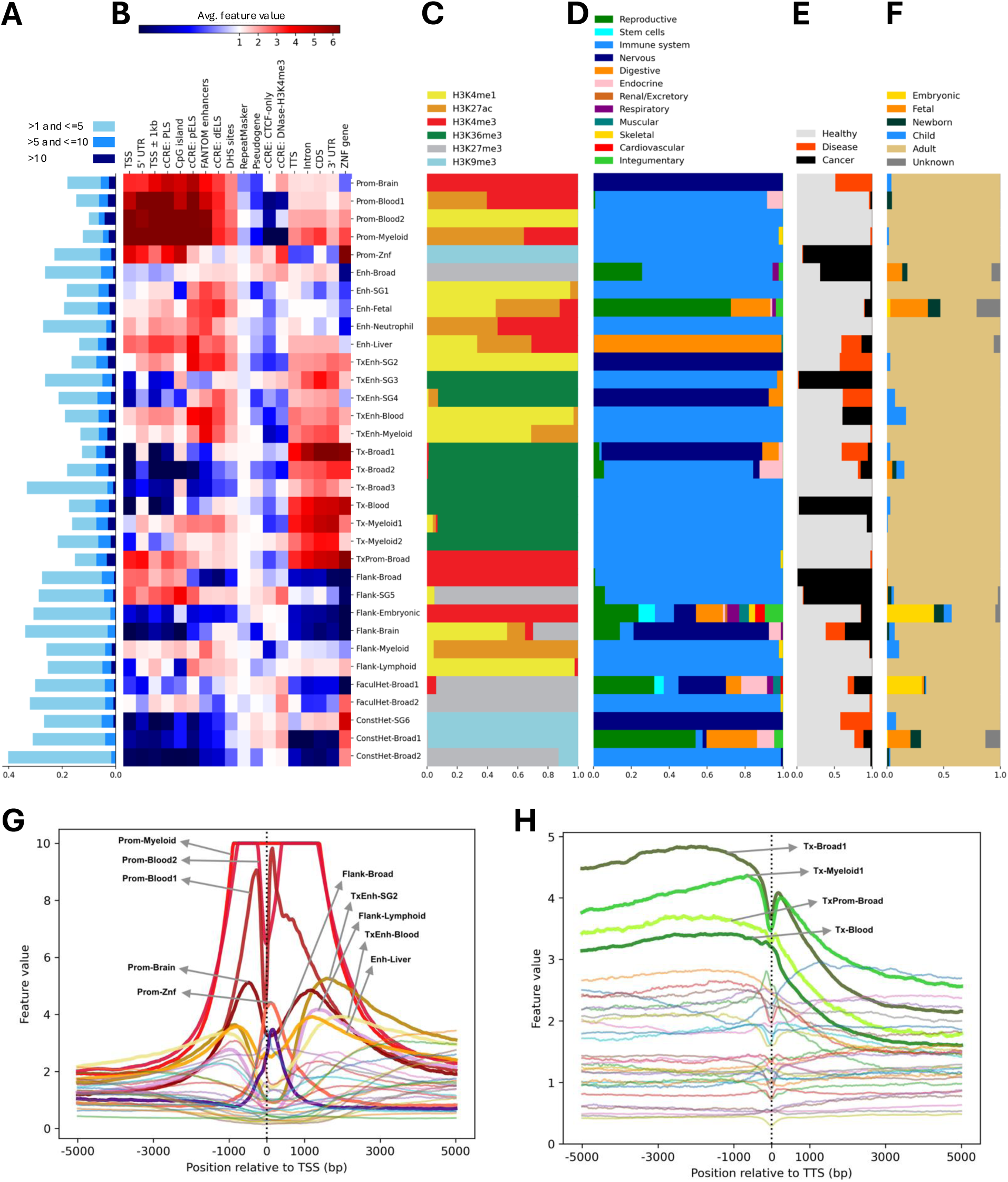
Pan-cell type features capture different histone modification marks and show distinct patterns around annotated genomic elements. **A** Distribution of feature values. Horizontal axis indicates the fraction of the genome with features in a given value range. Features are normalized to a genome-wide average of 1, so a value of 10 indicates 10 times the genome-wide average. **B** Average value of pan-cell type features at existing genome annotations. **C-F** Of the top 1% emission values, the histone modification (C), high-level cell type ontology (D), disease state (E), and donor life stage (F). **G-H** Average value of pan-cell type features around transcription start sites (**G**) and transcription termination sites (**H**) of protein-coding genes.

**Table 1:** Feature mnemonics. We assigned a mnemonic to each feature that explains the types of activity and cell types that it captures (section 5.10). In the third column, for brevity, we only indicate disease states when a feature is not exclusively associated with “Healthy” samples.

| Feature | Main emitted assays | Main emitted biosamples & disease states | Distribution |
| --- | --- | --- | --- |
| Prom-Brain | H3K4me3 | Nervous (Healthy & Disease) | Punctate |
| Prom-Blood1 | H3K4me3, H3K27ac | T-cell, Endocrine | Punctate |
| Prom-Blood2 | H3K4me3, H3K4me1 | Myeloid, other blood cell types | Punctate |
| Prom-Myeloid | H3K4me3, H3K27ac | Immune system | Punctate |
| Prom-Znf | H3K9me3, H3K4me3 | Immune system (Cancer) | Diffuse |
| Enh-Broad | H3K4me1, H3K27me3 | Reproductive, Immune system (Cancer), Respiratory, Integumentary | Diffuse |
| Enh-SG1 | H3K4me1, H3K27ac | Immune system, Liver | Punctate |
| Enh-Fetal | H3K4me1, H3K27ac | Reproductive, Digestive, Integumentary, Endocrine | Punctate |
| Enh-Neutrophil | H3K27ac, H3K4me3 | Immune system | Diffuse |
| Enh-Liver | H3K4me1, H3K27ac, H3K4me3 | Digestive (Healthy & Disease) | Punctate |
| TxEnh-SG2 | H3K4me1 | Nervous (Healthy & Disease), Immune system | Punctate |
| TxEnh-SG3 | H3K36me3 | Immune system (Cancer), Digestive (Healthy & Cancer) | Diffuse |
| TxEnh-SG4 | H3K36me3, H3K27ac | Nervous (Healthy & Disease), Digestive (Healthy & Disease) | Punctate |
| TxEnh-Blood | H3K4me1, H3K27ac | Immune system (Healthy & Cancer) | Punctate |
| TxEnh-Myeloid | H3K4me1, H3K27ac | Immune system | Punctate |
| Tx-Broad1 | H3K36me3, H3K4me3, H3K27ac | Nervous (Healthy & Disease), Digestive, Reproductive, Endocrine, Immune system | Punctate |
| Tx-Broad2 | H3K36me3 | Immune system, Endocrine (Healthy & Cancer), Reproductive, Nervous (Healthy & Cancer) | Punctate |
| Tx-Broad3 | H3K36me3, H3K4me3 | Broad | Diffuse |
| Tx-Blood | H3K36me3 | Immune system (Cancer) | Punctate |
| Tx-Myeloid1 | H3K36me3, H3K27ac | Myeloid | Punctate |
| Tx-Myeloid2 | H3K36me3, H3K27me3 | Myeloid | Diffuse |
| TxProm-Broad | H3K4me3, H3K36me3 | Immune system, Skeletal | Punctate |
| Flank-Broad | H3K4me3, H3K27ac | Immune system (Cancer), Reproductive, Endocrine, Digestive | Diffuse |
| Flank-SG5 | H3K4me1, H3K27ac, H3K27me3 | Immune system (Cancer), Reproductive | Diffuse |
| Flank-Embryonic | H3K4me3, H3K27me3 | Reproductive, Digestive, Nervous | Diffuse |
| Flank-Brain | H3K4me1, H3K27ac, H3K27me3 | Nervous (Healthy, Cancer & Disease), Reproductive | Diffuse |
| Flank-Myeloid | H3K4me1, H3K27ac | Immune system, Skeletal | Diffuse |
| Flank-Lymphoid | H3K4me1 | Immune system | Diffuse |
| FaculHet-Broad1 | H3K27me3 | Broad (Healthy & Cancer) | Diffuse |
| FaculHet-Broad2 | H3K27me3, H3K36me3, H3K9me3 | Broad | Diffuse |
| ConstHet-SG6 | H3K9me3 | Nervous (Healthy & Disease), Digestive | Diffuse |
| ConstHet-Broad1 | H3K9me3 | Broad | Diffuse |
| ConstHet-Broad2 | H3K9me3, H3K27me3 | Broad | Diffuse |

“Prom” features correspond to promoter activity, characterized by the marks H3K4me3 and H3K27ac in active cell types and H3K27me3 in inactive cell types (Fig. 3C, 4B, S5-S6, Table 1). As expected, Prom features are typically present at annotated TSSs (Fig. 3B and 3G). Additionally, we found that the spatial distribution of feature values varies depending on the type of epigenomic activities that each feature captures. For instance, Prom features tend to have a punctate distribution with zero or near zero values in most regions of the genome and high values (greater than 5) in the rest of the genome (Fig. 3A).

**Figure 4:**
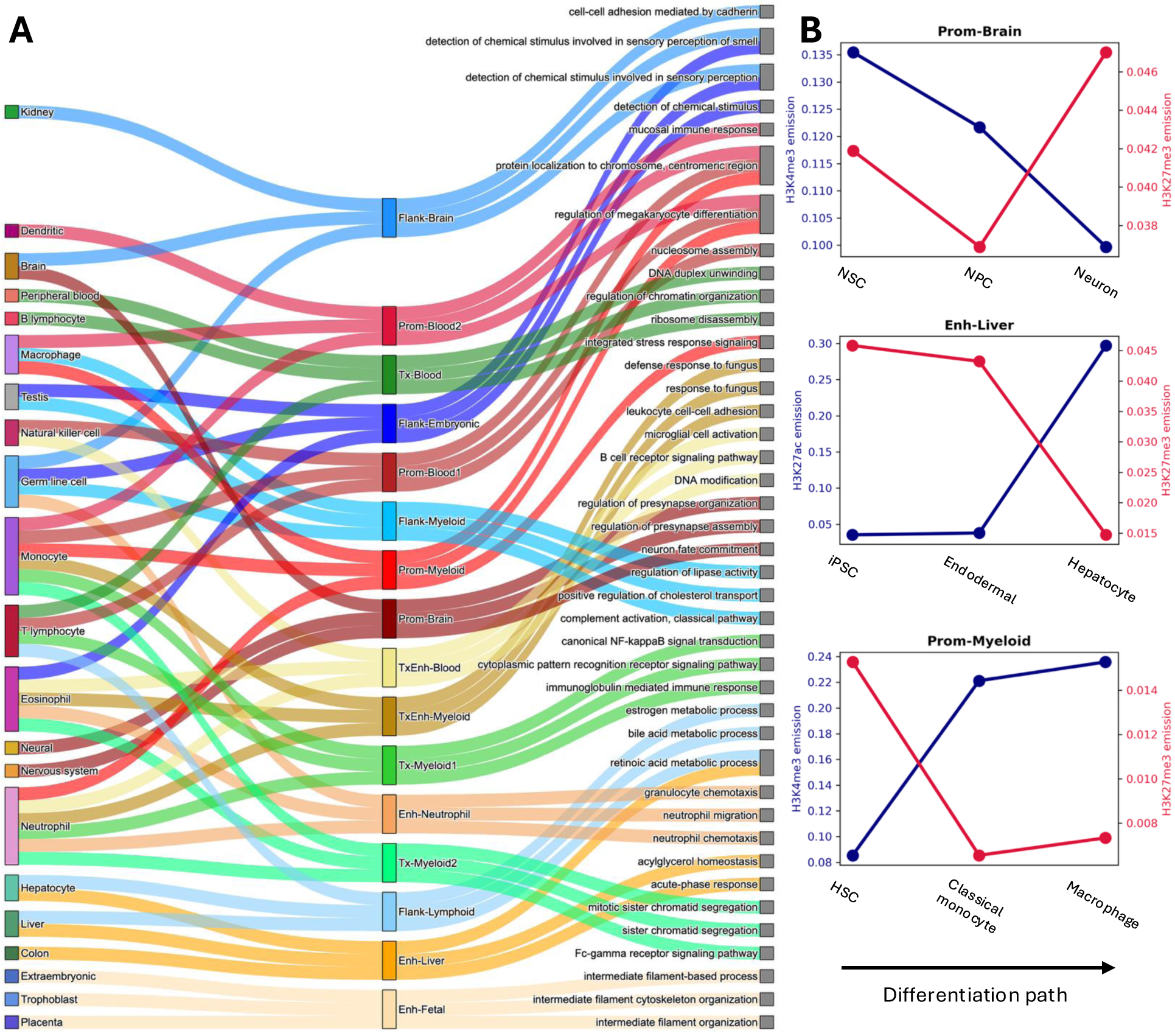
Pan-cell type features capture gene regulatory programs. **A** (left column) intermediate ontology; (middle column) pan-cell type features; (right column) Gene Ontology terms; The links connect each feature to the top three cell types from the gene expression analysis (section 5.8) and the top three GO terms from the gene ontology analysis (section 5.9). Only those features with a specific cell type/sample group are shown here. **B** Average emission of each feature for active marks vs. the repressive mark, H3K27me3, across several different stem and differentiated cells.

**Figure 5:**
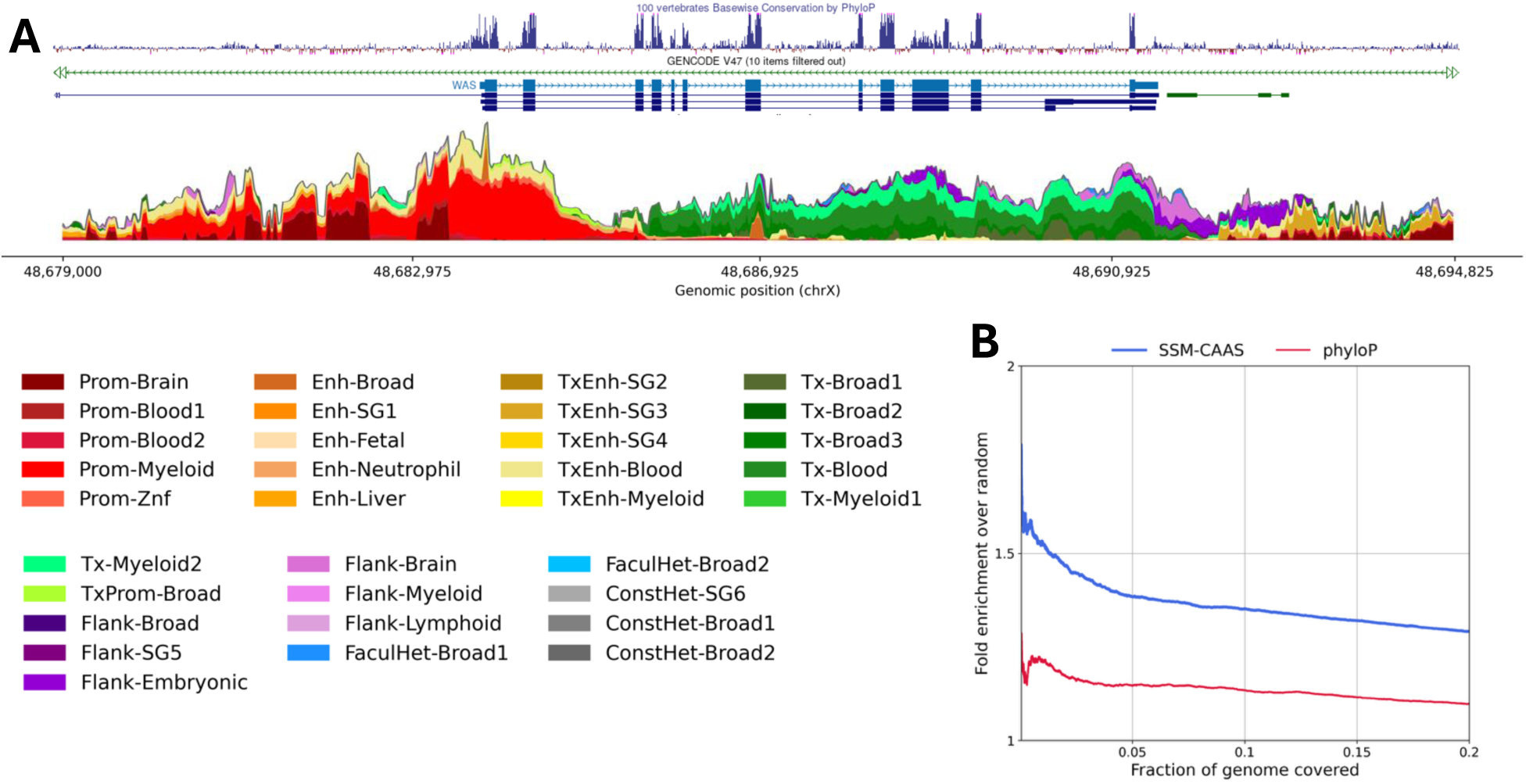
SSM-CAAS plot summarizes epigenomic signals at a given locus. **A** CAAS plot for the WAS locus. **B** Enrichment of SSM-CAAS and phyloP scores for GWAS variants.

“Enh” features correspond to enhancer activity, characterized by the marks H3K4me1 and H3K27ac in active cell types and H3K27me3 in inactive cell types (Fig. 3C, 4B, S4-S5, Table 1). These features are typically present at annotated enhancers and DHSs (Fig. 3B) and have a punctate distribution in these regions (Fig. 3A). “TxEnh” features correspond to enhancer activity of those enhancer elements that are typically found within the transcribed regions of the genome, such as the introns, characterized by the marks H3K4me1, H3K27ac, and H3K36me3 in active cell types and H3K27me3 in inactive cell types (Fig. 3C, S4-S5 and S7, Table 1).

“Tx” features are those that capture transcription activity, characterized by the mark H3K36me3 (Fig. 3C, S7, Table 1). As expected, Tx features are present at regions associated with transcription, such as the cod-ing regions (CDS), introns, and untranslated regions (UTRs) (Fig. 3B and 3H). Unlike the Prom and Enh features, Tx features tend to have a broad distribution, with moderate values (values 1-5) over broad regions (Fig. 3A).

Epigenetic patterns at the flanking regions to regulatory elements (e.g., promoters and enhancers) belong to a category we term “Flank” (Fig. 3B), characterized by a dispersed moderate signal in H3K4me1, H3K4me3, and H3K27ac (Fig. S4-S6, Table 1).

“FaculHet” and “ConstHet” refer to Faculatative and Constitutive heterochromatin, characterized by H3K27me3 and H3K9me3, respectively. We identified two ConstHet features capturing broad H3K9me3 signal across all samples, and another ConstHet feature capturing H3K9me3 in a subset of the samples (Fig. 3C, Table 1). In contrast, most H3K27me3 signal is captured by Prom and Enh features, which typically correspond to active regulatory marks in a few samples and H3K27me3 in others (Fig. S4-S7). As expected, heterochromatin features are depleted at annotated regulatory elements (Fig. 3B). Similar to the Tx features, both types of heterochromatin features tend to have a broad distribution, with moderate values (values 1-5) over broad regions.

Apart from these canonical activity types, we identified some other rarer types: Prom-Znf, characterized by H3K4me3 at zinc-finger genes (Fig. 3B-C, Table 1). Also, we have a “TxProm” feature that is present in both promoter and transcribed regions, characterized by the marks H3K4me3 and H3K36me3 (Fig. 3B-C, Table 1).

### 2.4 Pan-cell type features capture cell type-specific regulatory programs

We hypothesize that each feature corresponds to a specific regulatory program in the genome. Because many of the epigenomic activities underlying these programs co-vary across cell types, a relatively small set of 33 pan–cell type features can capture the major axes of variation across 9,539 tracks. As discussed, we found a few features correspond to broad activity, such as ConstHet-Broad1 and ConstHet-Broad2, corresponding to constitutive heterochromatin that is repressed in all samples, and Tx-Broad 1 to 3 corresponding to genes marked by H3K36me3 in all cell types which may play housekeeping roles.

However, such broad features are the minority. Most features learned by epigenome-ssm correspond to activity that is specific to a certain developmental lineage (Fig. 4A). For example, Prom-Brain captures promoter activities in brain-related cell types, while Enh-Liver captures enhancers active in livers (Fig. 3C-D, S8).

As expected, we found that genes that are expressed in a given cell type (according to RNA-seq data not used by epigenome-ssm) tend to have features representing active activity for that cell type (Fig. 4A, see Sec. 5.8). We found that for example, the Prom-Brain feature occurs at genes expressed in Brain, Neural, and Nervous system, while Prom-Myeloid occurs at genes expressed in Macrophage, Monocyte, and Neutrophil. This is consistent with the emissions for these two features, where the top emitted biosamples for Prom-Brain and Prom-Myeloid are Nervous and Immune system, respectively (Fig. 3C-D, see Fig. S8 for intermediate ontology).

We found that regulatory programs typically involve loci switching from active to repressed—or vice versa—throughout development. We found several cases where a feature captures the cell differentiation trajectory of a particular cell type group, starting with stem cells and ending with some mature differentiated cells (Fig. 4B). For instance, Prom-Brain emission for the active mark H3K4me3 decreases when moving from neural stem cells (NSCs) to neural progenitor cells (NPCs) and mature neurons, while the emission of this feature for the repressive mark H4K27me3 is at its highest at the end of the differentiation path for mature neurons. Similarly, Enh-Liver captures the differentiation path of iPSCs to endodermal and Hepatocyte cells, while Prom-Myeloid captures the differentiation path of HSCs to monocytes and macrophages.

Furthermore, by comparing features to Gene Ontology (GO) terms, we found that regulatory programs align with cellular functions (Fig. 4A). GO terms provide insights into the downstream functions of the genes in a hierarchical manner, starting with more general terms that correspond to functions carried out by most genes to more specific terms corresponding to functions that are carried out by a small subset of the genes. We assigned GO terms to each feature according to the cellular functions of genes this feature is associated with (Methods section 5.9). The GO terms assigned to each feature are highly relevant in the cell type/sample group of that feature. For instance, the top three GO terms for Prom-Brain are “neuron fate commitment”, “regulation of presynapse assembly”, and “regulation of presynapse organization” (Fig. 4A). These results provide evidence that patterns of regulatory activity correspond to patterns of gene regulation and cellular function.

### 2.5 Conservation-associated activity score (SSM-CAAS) captures activity relevant to phenotype and disease

The pan-cell type features generated by epigenome-ssm summarize activity across all cell types and tissues, and thus can identify activity important to organism-level phenotype and disease. It is established through genome-wide association studies (GWAS) that the majority of the disease-associated variants are in the non-coding regions of the genome, likely regions that contain some regulatory elements like enhancers and promoters; integrative annotations like epigenome-ssm are needed to identify these elements.

To enable this identification, we define SSM-CAAS, the SSM-based conservation-associated activity score (CAAS), based on the previously-defined SAGA-based CAAS [33]. Briefly, we used our pan-cell type features as input to a non-negative linear regression model that predicts evolutionary conservation (measured by phyloP [34]) and defined SSM-CAAS as the predicted phyloP score by this model. Evolutionary conservation is widely used as a proxy for putatively functional regions of the genome; thus SSM-CAAS identifies positions with regulatory activity consistent with putative function.

As expected, the contribution to SSM-CAAS differs between features (Fig. S9C). Features representing transcription have a high contribution to SSM-CAAS, as coding exons are generally the most conserved positions in the genome. Regulatory activity has SSM-CAAS that is comparable to transcription because, although these regions are typically less conserved than exons, epigenomic marks do not usually distinguish exons from less-conserved introns. As expected, features representing repressive or extremely cell type-specific activity have the lowest SSM-CAAS contribution.

The conservation-associated activity score offers two advantages over conservation as a measure of the importance of a given locus. First, the conservation-associated activity score is directly attributable to a specific activity in a specific set of cell types; in contrast, conservation indicates only that a position is important, with no way to determine how it acts. SSM-CAAS has the potential to mark elements that are not conserved but have epigenetic activity similar to other conserved elements, such as those that gained regulatory or transcriptional activity only in humans. Conversely, SSM-CAAS can identify elements conserved across other species but decommissioned in humans.

Furthermore, SSM-CAAS enables a visualization of genomic activity that emphasizes putative function. This visualization consists of a stacked ribbon plot in which the total height of the plot is the predicted conservation of that locus, and the colors of the ribbon constitute the contribution of each activity type to the prediction (Fig. 5A, S9A-B). We found that positions with high SSM-CAAS are more enriched for genetic variants identified by genome-wide association studies (GWAS) to be involved in disease (Fig. 5B), compared to phyloP. This indicates that CAAS captures regions of the genome that are not only evolutionarily conserved but also functionally relevant to human disease, providing a valuable tool for identifying disease-associated regulatory elements.

Several examples show how epigenome-ssm features can be used to quickly understand a genomic locus. By diving deep into several different genomic loci, we found that SSM-CAAS not only captures the main genomic elements (e.g., promoters, transcribed genes, etc.), but it also captures cell type-specific activities (Fig. 5A, S9A-B). The WAS gene located on the forward strand (Fig. 5A) has a conserved promoter region, as indicated by the phyloP track, and has a high SSM-CAAS signal that is mainly derived from promoter features (in reds). The dominant promoter feature for WAS is Prom-Myeloid, and that is expected given the fact that WAS is highly expressed in the immune cells [35]. The transcription-associated features (in greens) cover the gene body while there are enhancer features (in yellows) upstream of the promoter, within the introns, and downstream of the gene.

The ADPGK gene is located on the reverse strand (Fig. S9A). Similar to WAS, the promoter features cover the region around the transcription state site while the Tx features cover the gene body. This gene also has some enhancers as indicated by the Enh and TxEnh features (yellows). The CXCR5 gene located on the forward strand is surrounded by a few other genes (Fig. S9B). The promoter features are enriched around the promoter region of CXCR5, and also the promoter regions of both the upstream and downstream genes. While the enhancer features are very high around CXCR5 (likely due to the proximal enhancers regulating the downstream genes like BCL9L), the TxBroad features are enriched for the transcribed regions of the downstream gene, BCL9L.

These examples show how visualizing epigenome-ssm features and SSM-CAAS can allow a user to quickly understand the activity of a given locus across all 1,698 biosamples, both in terms of types of activity and tissue specificity.

## 3 Discussion

In this study, we introduced a continuous pan-cell type chromatin state annotation and demonstrated its effectiveness in summarizing the vast epigenomic data from IHEC. Our results showed that these pan-cell type features are capable of capturing biologically meaningful patterns in a variety of genomic contexts. The ability of epigenome-ssm to provide a more nuanced representation of chromatin state activity across multiple cell types has significant implications for understanding gene regulation and disease association.

One key advantage of continuous annotations over discrete ones is their ability to capture varying levels of chromatin activity at each genomic position, allowing for a more accurate representation of the regulatory landscape. As demonstrated in our comparisons, even with fewer features (K=16), epigenome-ssm matched or exceeded the performance of ChromHMM, which used 100 discrete labels. This suggests that the continuous nature of our features allows for more efficient encoding of regulatory information, potentially reducing redundancy while still capturing the complexity of the underlying epigenetic signals.

Furthermore, the cell type-specific patterns identified by epigenome-ssm emphasize its utility in studying tissue-specific regulatory programs. Features such as Prom-Brain, Enh-Liver, and Prom-Myeloid demonstrated strong associations with their respective cell type groups, suggesting that these features effectively capture regulatory programs that drive tissue-specific gene expression. This ability to distinguish between cell type-specific activities is crucial for applications in disease association studies, as it allows researchers to pinpoint regulatory elements that may be disrupted in specific tissues or cell types.

Beyond the canonical promoter, enhancer, transcription, and heterochromatin patterns, our model recovers rarer mixed signatures that likely reflect specialized regulatory contexts. First, Prom-Znf shows strong promoter-like H3K4me3 at zinc-finger genes against a broader H3K9me3 background, consistent with the known organization of ZNF clusters (e.g., in chr19) in constitutive/constitutive-like heterochromatin while maintaining promoter activity at specific loci; such coupling may reflect tight transcriptional control with context-dependent activation. Second, TxProm exhibits concurrent promoter (H3K4me3) and transcriptional (H3K36me3) marks at the same loci, compatible with overlapping/alternative TSS usage within transcribed regions. These non-canonical states illustrate how continuous features capture combinatorial activities.

Finally, our CAAS analysis provides further evidence of the utility of pan-cell type features in functional genomics. The enrichment of disease-associated variants within regions with high CAAS values suggests that our method can prioritize genomic regions of interest for further investigation, particularly in the context of non-coding disease-associated variants identified by GWAS. This is particularly important given the increasing recognition that many disease-associated variants lie outside of coding regions, in regulatory elements that control gene expression.

We expect that annotations from epigenome-ssm will provide an easily-interpretable summary of epige-netic activity across hundres of cell types and tissues, and thus it will be the first stop for researchers aiming to understand the epigenome.

## 4 Methods

### 4.1 Input data and processing

All the analyses are done using the human reference genome hg38. From the IHEC repository, we downloaded 9,539 bigWig tracks containing − log_10_ *p-value* signals from six core histone modification ChIP-seq assays. Those tracks contain imputed ChIP-seq signals and were generated using ChromImpute [36] at 25 bp resolution from 5,339 observed ChIP-seq tracks in the IHEC repository. To reduce the effect of outliers, we transformed the signals using the inverse hyperbolic sine function defined as *arcsinh*(*x*) = log(*x* + √*x*^2^ + 1).

We trained epigenome-ssm on the ENCODE Pilot regions [37], which consist of 1% of the human genome, and used the UCSC liftOver tool [38] to convert the ENCODE Pilot coordinates from hg19 to hg38. We also removed the ENCODE exclusion list regions [39] from all the analyses. We then used the trained epigenome-ssm model to generate annotations for the whole genome.

### 4.2 epigenome-ssm model, training procedure, and annotation pipeline

The epigenome-ssm model is based on the state-space model (SSM) and is an unsupervised linear sequential model that takes as input *E* observed epigenomic signal values for each position *g* of the genome and generates *K* non-negative chromatin state feature values that summarize epigenomic activity at position *g*. More specifically, this model has two main assumptions: (1) At each position *g*, there is a latent vector *α_g_* ∈ R*^K^* that encodes the chromatin state features at *g* and the observed data *y_g_* ∈ R*^E^* is generated as a linear function of *α_g_*, parameterized by an emission matrix *Z* ∈ R*^E×K^*; and (2) *α_g_*_+1_ is generated as a linear function of *α_g_*, parameterized by a transition matrix *T* ∈ R*^K×K^*. To limit the model’s capacity to overfit and its sensitivity to local optima, we additionally added two *L*_2_ regularization terms to the optimization’s objective function *J* (*Z, T*), which encourage *Z* and *T* to have small values: *J* (*Z, T*) = log *P* (*α, Z, T* |*Y*) + *λ*_1_∥*Z*∥*_F_* + *λ*_2_∥*T* ∥*_F_*. Additionally, we used an active set method of Lagrange multipliers to enforce non-negativity constraints on both chromatin state features and the emission parameters (see [30] for more details).

To train the model, we used an expectation maximization (EM) algorithm to maximize the model’s log-likelihood as a function of its parameters, *Z* and *T* . We generated ten random initializations of the model parameters and took the one with the best log-likelihood. We did not use a fixed number of itera-tions—instead, we kept the training going until there was no significant improvement in log-likelihood.

As a normalization step at the end of the pipeline, we normalized each feature *i* such that its genome-wide mean is 1, and correspondingly multiplied its emission values *Z*_:*,i*_ by its unnormalized genome-wide mean. This step leaves the predicted observed data unchanged, but makes it easier to interpret the features because they are all on the same scale.

### 4.3 Alternative pan-cell type annotations

In our quantitative evaluations, we included pan-cell type annotations generated by four other methods: DHS-NMF [28], Sei [31], Universal ChromHMM [29], and CSREP [32].

DHS-NMF annotations were generated using DNase-seq data across 733 human biosamples. The DNase-seq assay measures chromatin accessibility at DNase I hypersensitive sites (DHSs). More specifically, the authors curated a set of approximately 3.6 million DHSs across all the samples and then ran non-negative matrix factorization with 16 components on a DHS-by-biosample binary matrix. Therefore, DHS-NMF annotations are only available at DHSs and not the whole genome. We used both the DHS coordinates [40] and DHS-NMF annotations [41] in our analyses.

Sei annotations were generated by clustering the predictions from the Sei model, which is a deep learning model that predicts epigenomic signals from DNA sequence [31]. We downloaded a BED file from [42] that contains annotation of genomic positions with Sei’s sequence classes.

From the IHEC repository, we downloaded Universal ChromHMM annotations generated by Ernst Lab at UCLA by running ChromHMM on six core histone modification ChIP-seq data across all IHEC epigenomes. This set of annotations is generated using a model with 100 labels. Note that these annotations are similar, but not identical, to annotations presented in [29] that were generated using a smaller data set taken from only Roadmap Epigenomics.

We also downloaded the CSREP annotations generated by Ernst lab from the IHEC repository. The annotations were initially generated using a ChromHMM model with 18 states. Specifically, for the discrete version (termed as ”CSREP-dis in our evaluations”), we downloaded 56 BED files corresponding to the summarized annotations of 56 sample groups in the IHEC dataset. For the continuous version (i.e., CSREP-con), we downloaded 1,008 BigWig files containing CSREP probabilities for 18 labels across 56 sample groups.

### 4.4 Gene expression prediction

We evaluated the pan-cell type annotations based on their association with gene expression, as measured by bulk RNA-seq. We downloaded RNA-seq TPM (transcripts per kilobase million) data for 1,467 epigenomes from the IHEC repository. We then compared epigenome-ssm, Universal ChromHMM, CSREP, Sei, and DHS-NMF by using annotations within the gene body (i.e., the region between the transcription start site and transcription termination site) as predictors in a linear regression model to predict RNA-seq TPM values. More specifically, for a given epigenome-ssm model, we constructed a vector of *K* values from a given gene’s body by averaging each feature in that region, where *K* is the number of features for each model. For ChromHMM, we formed a one-hot encoded vector of length 100 corresponding to the 100 labels in the Universal ChromHMM annotation. Position *i* of this vector has a value of 1 if label *i* is present in the region, and 0 otherwise. To account for the length of each label in the region, we multiplied each value of the vector by the proportion of the region covered by the corresponding label. Similarly, we formed a one-hot encoded vector of length 18 for each of the 56 sample groups in CSREP annotation set. For the continuous version of the CSREP annotation (i.e., CSREP-con), we used the CSREP probability values instead of the discrete labels. For Sei, similar to ChromHMM, we formed a one-hot encoded vector of length 40 corresponding to Sei’s 40 sequence classes. For DHS-NMF, we formed a vector of length 16 corresponding to the 16 NMF components by averaging the value of each component across DHSs within each gene’s body. As the regression response value, we formed a vector of length 1,467 for each gene, corresponding to its RNA-seq TPM values across all epigenomes. We *arcsinh* transformed both predictor and response values (section 5.1).

We performed a similar analysis with all the same steps as above, except that we used annotations within the promoter region instead of the gene body. Specifically, for each gene, we centered a 2 kb window at the annotated TSS and extracted the annotations from that window.

We generated linear regression predictions in a 10-fold cross-validation setting and used Pearson and Spearman correlations to measure prediction performance. We limited this analysis to protein-coding genes and used GENCODE v45 [43] gene annotations. Also, we used the *LinearRegression* model from scikit-learn [44] for this analysis, as well as for the prediction tasks in sections 5.5 and 5.6.

### 4.5 Enhancer activity prediction

We evaluated the pan-cell type annotations based on their association with enhancer activity, as measured by Cap Analysis of Gene Expression (CAGE). We downloaded CAGE TPM data for 1,827 samples [45] and enhancer annotations [46] from FANTOM5 and used the UCSC liftOver tool [38] to convert the coordinates from hg19 to hg38. As the regression predictor value, we extracted annotations that overlap enhancers for each model, as described in section 5.4. As the regression response value, we formed a vector of length 1,827 for each enhancer, corresponding to its CAGE TPM values across all samples. We *arcsinh* transformed both predictor and response values (section 5.1).

### 4.6 Evolutionary conservation prediction

We evaluated the pan-cell type annotations based on their association with evolutionary conservation, as measured by phyloP [34]. We limited this analysis to the DHSs since DHS-NMF annotations are only available for those regions. We downloaded a phyloP track from the UCSC Genome Browser that is generated by multiple alignments of 99 vertebrate genomes to the human genome [47]. As the regression predictor value, we extracted annotations that overlap DHSs for each model, as described in section 5.4. As the regression response value, we used the absolute value of the mean phyloP score for each DHS. We *arcsinh* transformed both predictor and response values (section 5.1).

### 4.7 Enrichment of pan-cell type features for annotated genomic elements

We computed the enrichment of a pan-cell type feature for a given set of genomic regions as the mean feature value at those regions divided by the feature’s genome-wide mean. The following annotations were used:

- We used GENCODE v45 [43] for all gene annotations and only included protein-coding genes. We took the start position of the genes as TSS and the end position of the genes as TTS. “TSS ± 1kb” includes 2 kb regions centered at TSSs. Coding sequence (CDS) regions correspond to the GENCODE elements whose feature type is “CDS”. The 5’ and 3’ UTRs (untranslated region) correspond to the GENCODE elements whose feature type is “five prime UTR” “three prime UTR”, respectively. Similarly, exons correspond to the GENCODE elements whose feature type is “exon”. For introns, we took the elements whose feature type is “transcript” and removed the parts that overlapped with exons using the *bedtools subtract* command [48]. For zinc finger genes, we took those elements whose feature type is “gene” and their “gene name” contains “ZNF”. For pseudogenes, we took the elements with “gene” feature type and “pseudogene” in their “gene type” and included both protein- and non-protein-coding genes.
- We downloaded the ENCODE candidate cis-regulatory elements from the ENCODE’s SCREEN portal [49] and extracted the coordinates for the PLS (TSS-overlapping with promoter-like signatures), pELS (TSS-proximal with enhancer-like signatures), dELS (TSS-distal with enhancer-like signatures), DNase-H3K4me3 (not TSS-overlapping and with high DNase and H3K4me3 signals only), and CTCF-only (not TSS-overlapping and with high DNase and CTCF signals only) [50].
- The annotations for CpG islands [51] and RepeatMasker regions [52] were downloaded from the UCSC Genome Browser.
- Enhancer annotations were downloaded from FANTOM5 [46] (see section 5.5).
- We used the DHS coordinates [40] from [28] (see section 5.3).

### 4.8 Associating features with cell types through gene expression

We associated the pan-cell type features with cell type groups by looking at the feature values at protein-coding genes and the expression of those genes in different cell types. More specifically, we generated two matrices: (1) A feature-by-gene matrix that contains the average value of each feature at protein-coding genes;(2) A cell type-by-gene matrix that contains the RNA-seq expression value of each gene in different cell types. We generated the product of those two matrices over the gene axis to get a feature-by-cell type matrix. We downloaded RNA-seq data for 1,467 epigenomes from the IHEC repository for this analysis.

### 4.9 Associating features with Gene Ontology terms

We associated the pan-cell type features with gene ontology (GO) [53, 54] terms by looking at the feature values at protein-coding genes and the GO terms assigned to each gene in the GO database. More specifically, we generated two matrices: (1) A feature-by-gene matrix that contains the average value of each feature at protein-coding genes; (2) A GO term-by-gene binary matrix that connects each gene to the GO terms assigned to it. We generated the product of those two matrices over the gene axis to get a feature-by-GO term matrix. Since GO terms are assigned to each gene in a hierarchical manner (i.e., each gene may receive several GO terms starting from the most general term to more specific terms), we included all the terms assigned to each gene in our analysis except those terms that were assigned to less than 30 genes. For this analysis, we downloaded GO data files from [55, 56].

### 4.10 Assigning feature mnemonics

A feature mnemonic summarizes the primary activities captured by a pan–cell type feature. Because this study includes a large number of input signal tracks that can be characterized along multiple axes (e.g., histone modification assay, cell ontology, disease state, donor life stage and so on), we kept the mnemonics concise by including only the activity type and the cell ontology. We drew on multiple sources of evidence and examined each feature from several complementary perspectives. The materials used to assign mnemonics are summarized in Table 1, Fig. 3, Fig. 4, and Fig. S8. The specific steps are as follows.

For activity assignment, we combined (i) feature enrichment around public genome annotations (GENCODE, ENCODE, FANTOM, and DHS sites) with (ii) high emission values for the relevant histone modification assays. We considered a feature enriched for a given annotation if its average value across the annotated regions was ≥ 2. Because feature values are normalized by the genome-wide mean, a value of 2 indicates a twofold enrichment relative to the genome-wide average.

For cell type assignment, we integrated emission values, RNA-seq gene expression, and Gene Ontology (GO) to assign the cell type component of the mnemonic. For each feature, we ranked all input tracks by emission for that feature and computed the proportion of total emission contributed by each of the 12 high-level ontologies (Fig. 3D) and by each of the 40 intermediate ontologies (Fig. S8B). In general, we used the dominant high-level ontology in the mnemonic unless a dominant intermediate ontology was supported by both the gene expression and GO analyses (i.e., that intermediate ontology appears among the top three cell types in the expression analysis and its top GO terms are consistent; Fig. 4A). When several closely related cell types were dominant, we used a higher-order label (e.g., “Prom-Myeloid”: top emitters are Monocyte and Macrophage, both within the myeloid lineage). When the dominant cell types were not closely related, we used “SG” (Sample Group) in the mnemonic. Finally, we used “Broad” for features associated with a wide range of cell types that could not be cleanly grouped into a higher-order ontology.

### 4.11 Conservation-associated activity score

We generated a genome-wide track of conservation-associated activity score using both the pan-cell type features as a summary of the human epigenomic data across many cell types, and evolutionary conservation as measured by phyloP [34]. We used the same phyloP track as in section 5.6 and trained a non-negative linear regression model that takes as input the pan-cell type features and predicts the absolute phyloP score for each genomic bin. We performed a 5-fold cross validation and used the average coefficients as the final model to generate a predicted phyloP track. We defined CAAS to be the predicted phyloP scores from this model.

### 4.12 Enrichment for GWAS variants

We evaluated the conservation-associated activity score generated from our pan-cell type features based on the enrichment for disease-associated variants. We downloaded a set of disease-associated variants from GWAS Catalog [57]. We sorted the genomic bins based on their SSM-CAAS (and similarly based on their phyloP score), found the number of GWAS variants covered by each bin, and plotted the fold enrichment of SSM-CAAS and phyloP over the genome-wide average.

## 5 Code and data availability

Here is a link to a UCSC genome browser session containing the pan-cell type feature tracks along with the SSM-CAAS track: https://genome.ucsc.edu/s/habib/epigenome%2Dssm.IHEC.compact

Also, the latest code for the epigenome-ssm pipeline can be found here: https://github.com/habibdanesh/ epigenome-ssm

## Acknowledgements

We would like to thank the Ernst Lab at UCLA for generating ChromImpute imputations. We would also like to thank Guillaume Bourque, Stephan Beck, and other members of the IHEC community for their constructive feedbacks.

