## Supplementary figures for "Pan-cell type continuous chromatin state annotation of all epigenomes from the International Human Epigenome Consortium"

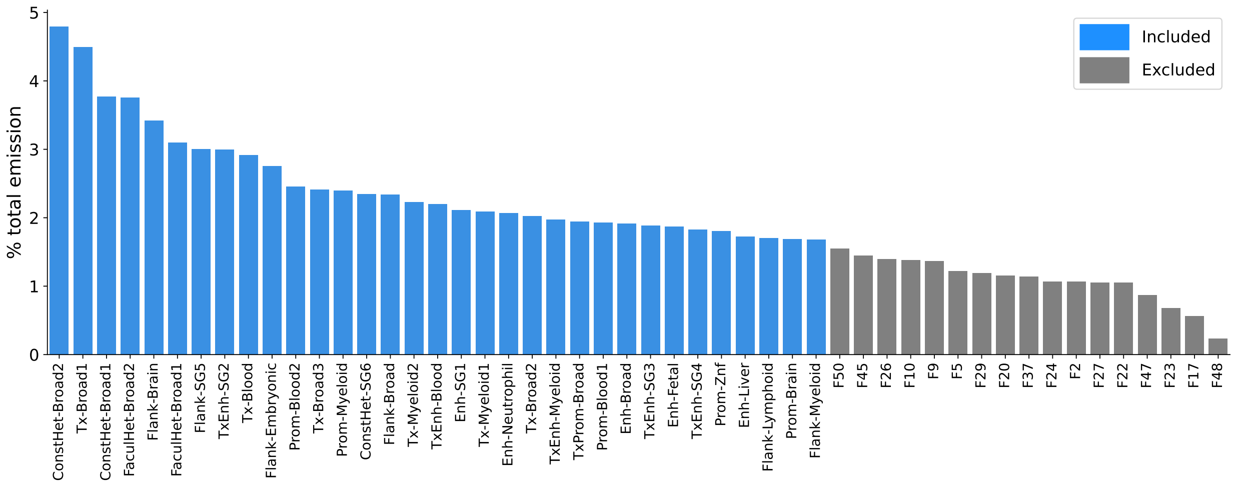


Figure S1: We trained a model with 50 features and took the 33 features with the highest total emission.

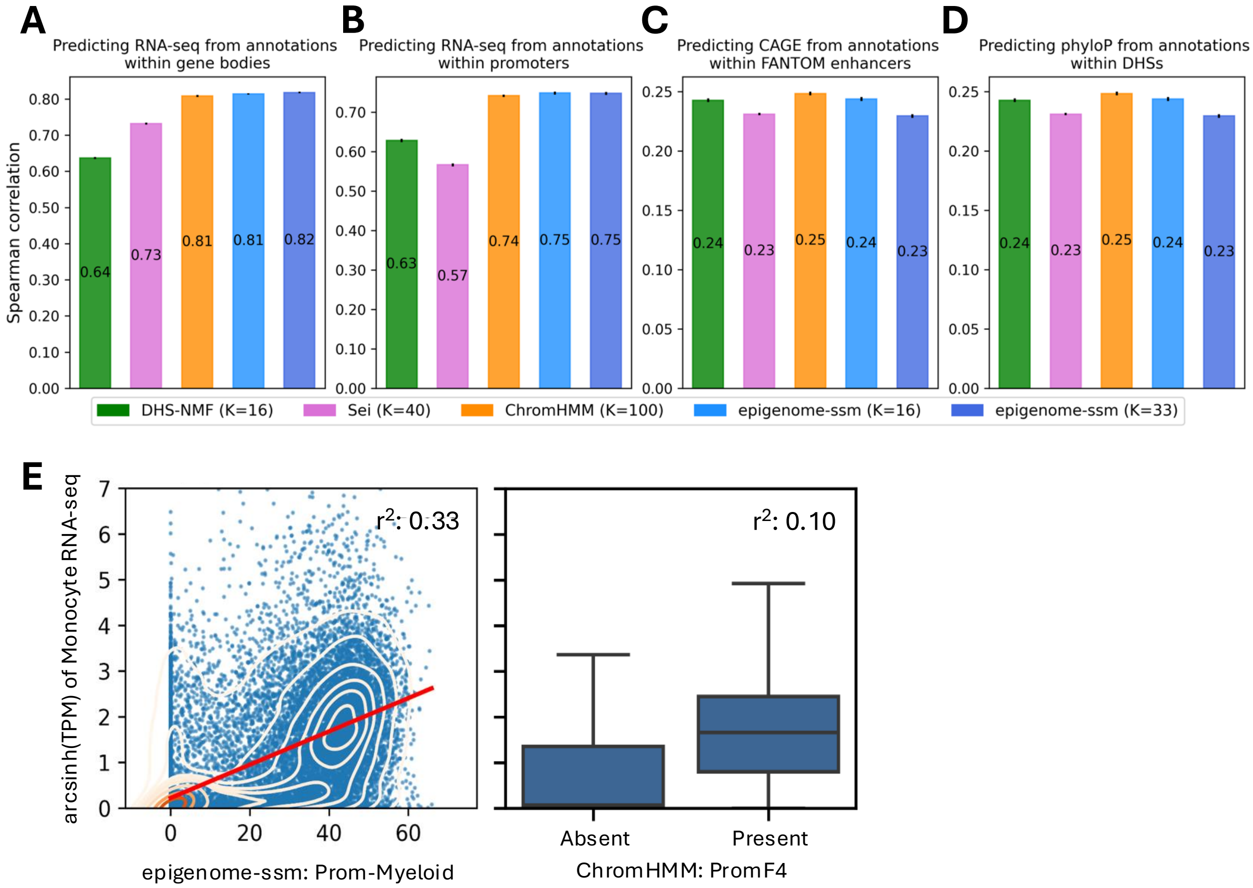


Figure S2: Pan-cell type features are predictive of genomic activities. (A-D) Evaluating pan-cell type annotations according to their prediction power of various genomic phenomena, similar to the plots in Fig. 2, but showing the Spearman correlation. $K$ is the number of components, sequence classes, labels, and features for DHS-NMF, Sei, ChromHMM, and epigenome-ssm, respectively. The vertical solid lines at the top of the bars represent the standard error from a 10-fold cross-validation. **A** Using annotations from within gene bodies to predict RNA-seq TPM values across all epigenomes. **B** Same as **A** but using annotations within a 2 kb window centered at TSS. **C** Using annotations from within FANTOM5 enhancers to predict CAGE TPM values across all samples. **D** Using annotations from within DHSs to predict evolutionary conservation, as measured by phyloP. **E** Predicting RNA-seq from a single feature/label (left) The epigenome-ssm feature with the highest correlation with Monocytes gene expression. The X-axis shows Prom-Myeloid feature value for each gene; (right) The ChromHMM label with the highest correlation with Monocytes gene expression. The box plots show the distribution of RNA-seq readouts for genes which are labeled with PromF4 (Present) and genes which are not labeled with PromF4 (Absent).

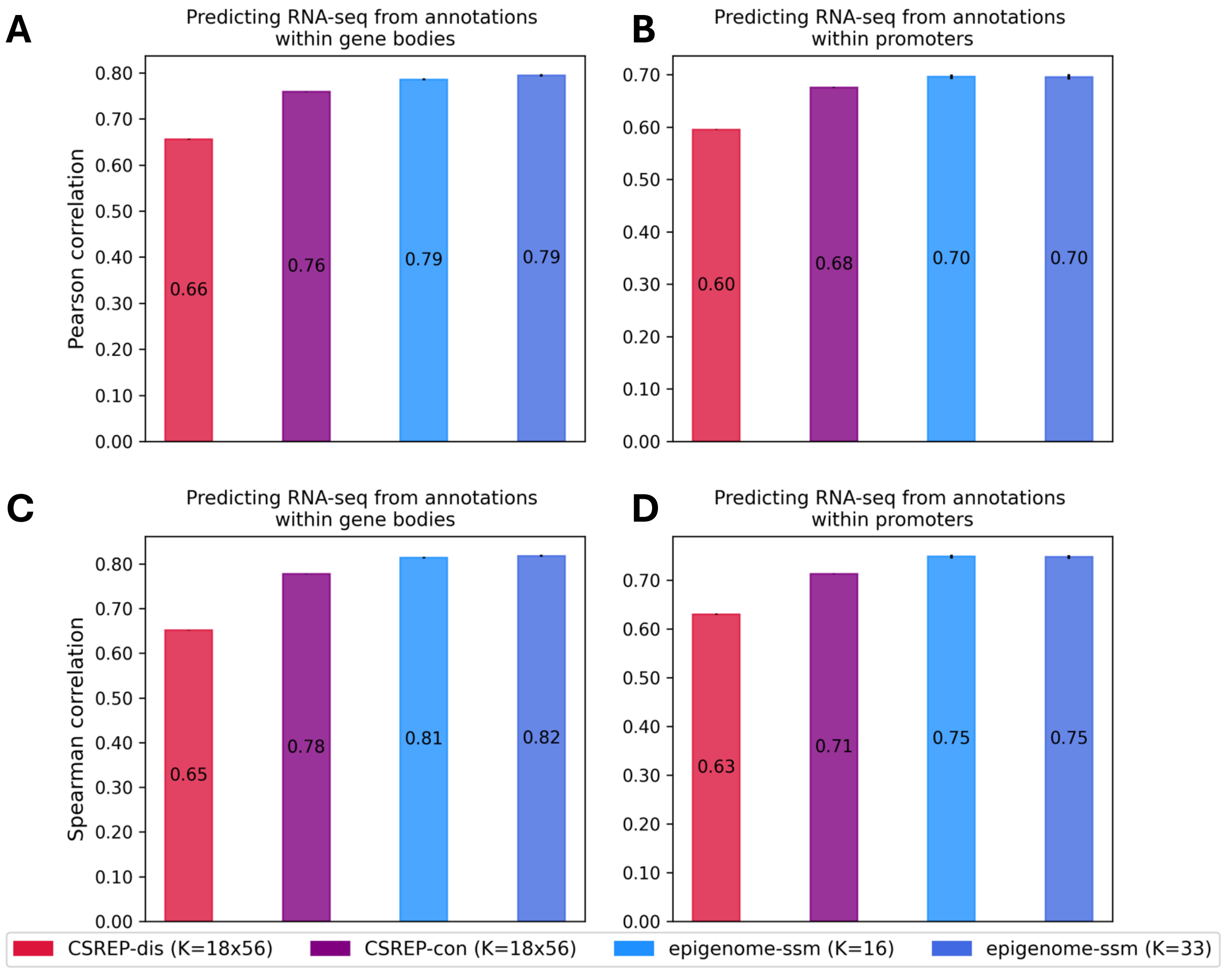


Figure S3: Pan-cell type features are predictive of genomic activities. (A-D) Evaluating pan-cell type annotations according to their prediction power of various genomic phenomena. $K$ is the number of labels and features for CSREP and epigenome-ssm, respectively. The vertical solid lines at the top of the bars represent the standard error from a 10-fold cross-validation. CSREP annotations are per-sample-group (56 in total) and are generated using a ChromHMM model with 18 labels. CSREP-dis refers to discrete labels and CSREP-con refers to continuous probability values. **A** Using annotations from within gene bodies to predict RNA-seq TPM values across all epigenomes. **B** Same as **A** but using annotations within a 2 kb window centered at TSS. **C** Same as **A** but showing Spearman correlation. **D** Same as **B** but showing Spearman correlation.

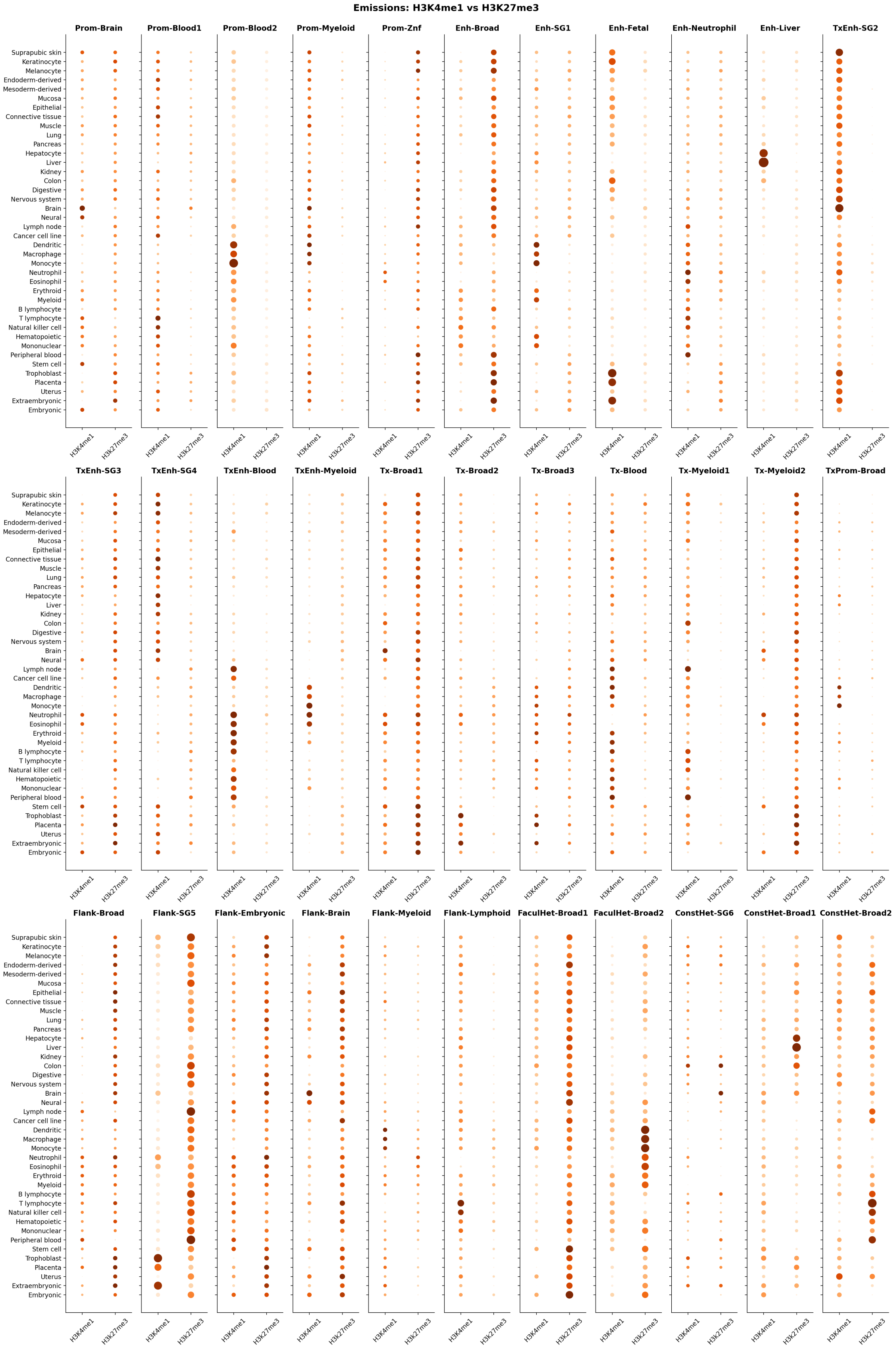


Figure S4: Emission values of the pan-cell type features for the active mark, H3K4me1, and the repressive mark, H3K27me3, for input samples represented by their intermediate ontology.

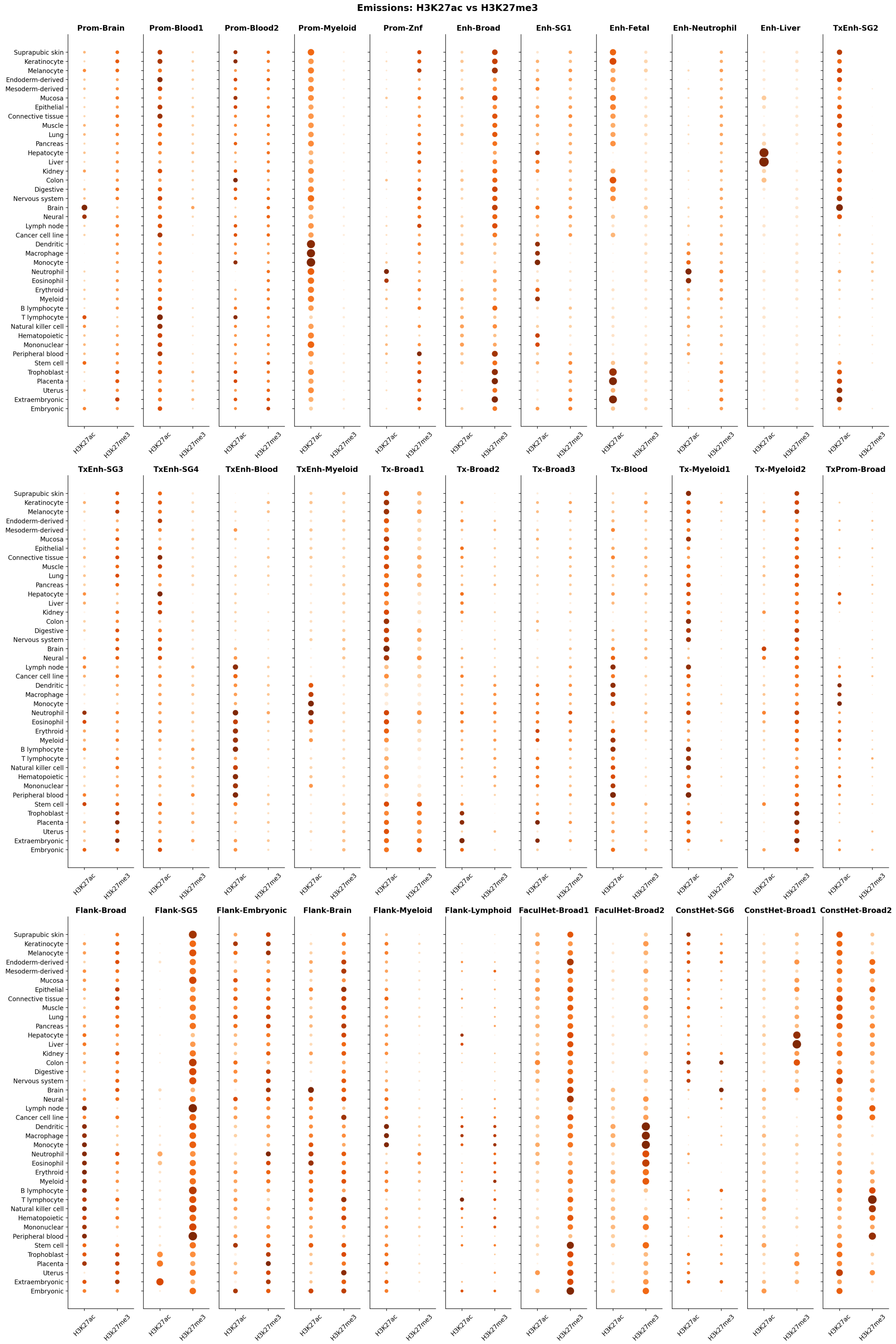


Figure S5: Emission values of the pan-cell type features for the active mark, H3K27ac, and the repressive mark, H3K27me3, for input samples represented by their intermediate ontology.

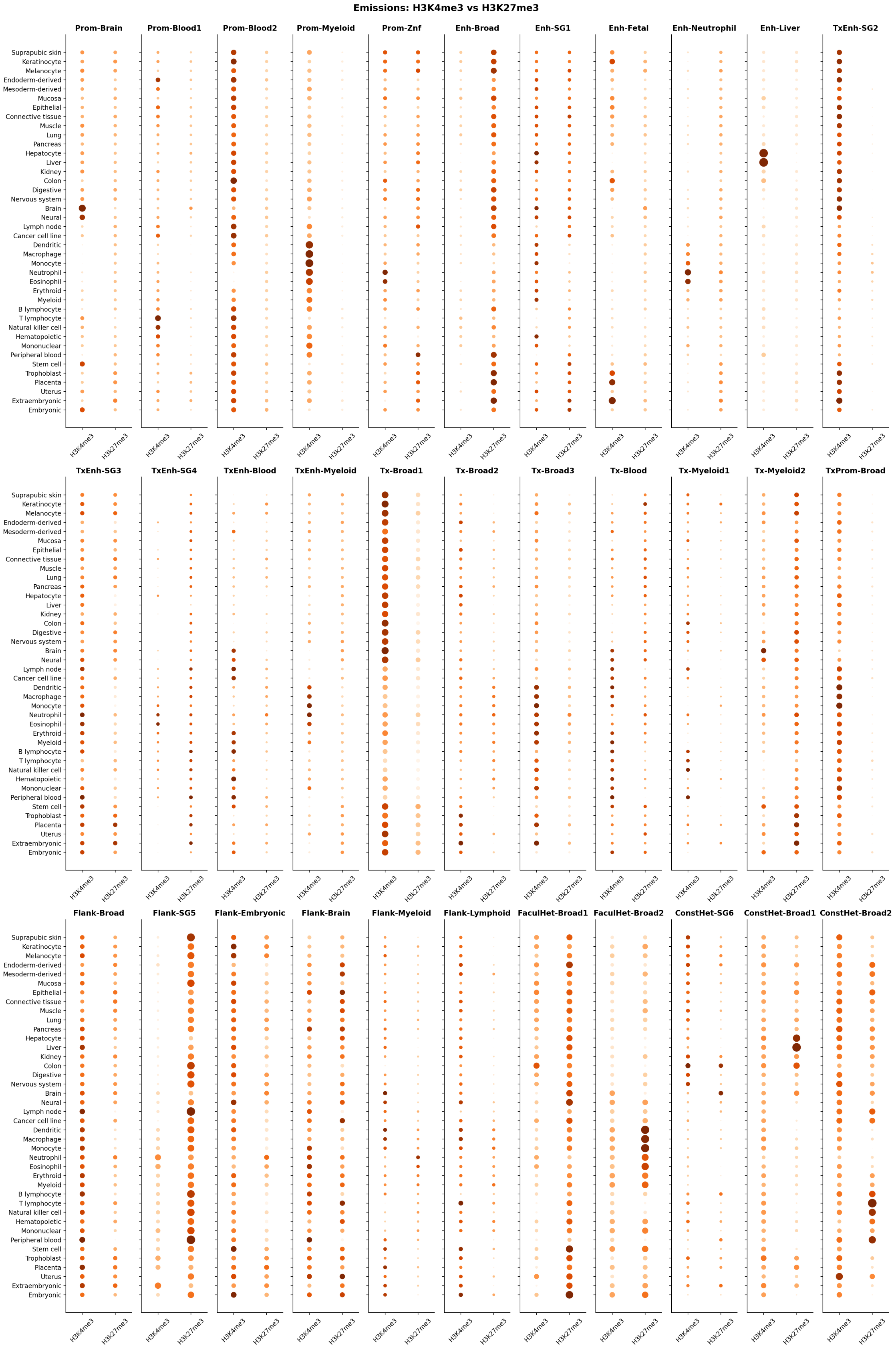

Figure S6: Emission values of the pan-cell type features for the active mark, H3K4me3, and the repressive mark, H3K27me3, for input samples represented by their intermediate ontology.

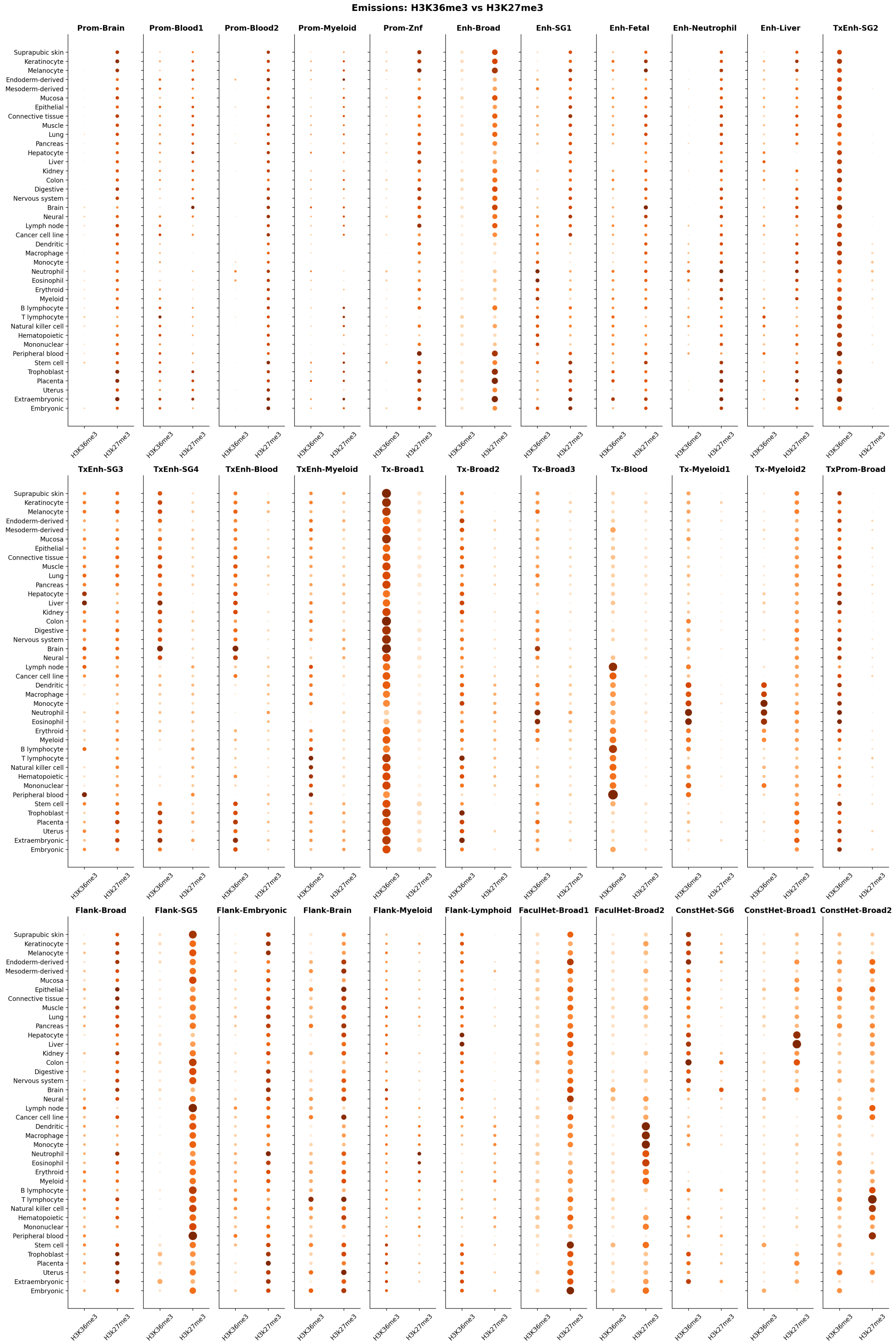


Figure S7: Emission values of the pan-cell type features for the active mark, H3K36me3, and the repressive mark, H3K27me3, for input samples represented by their intermediate ontology.

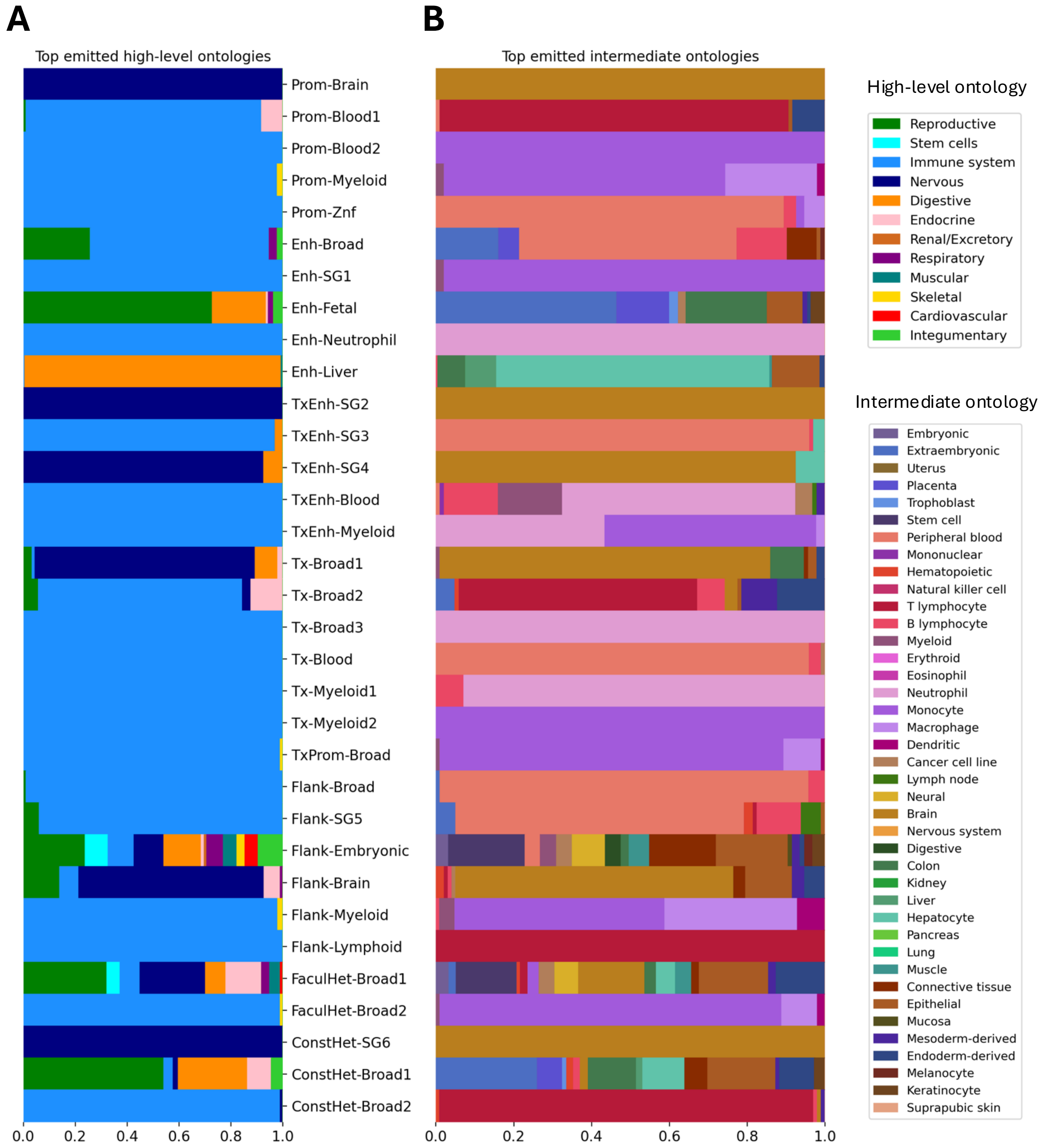


Figure S8: Portion of the total emission from the top 1% of input signal tracks for each pan-cell type feature. **A** Input tracks colored based on high-level ontology. **B** Input tracks colored based on intermediate ontology.

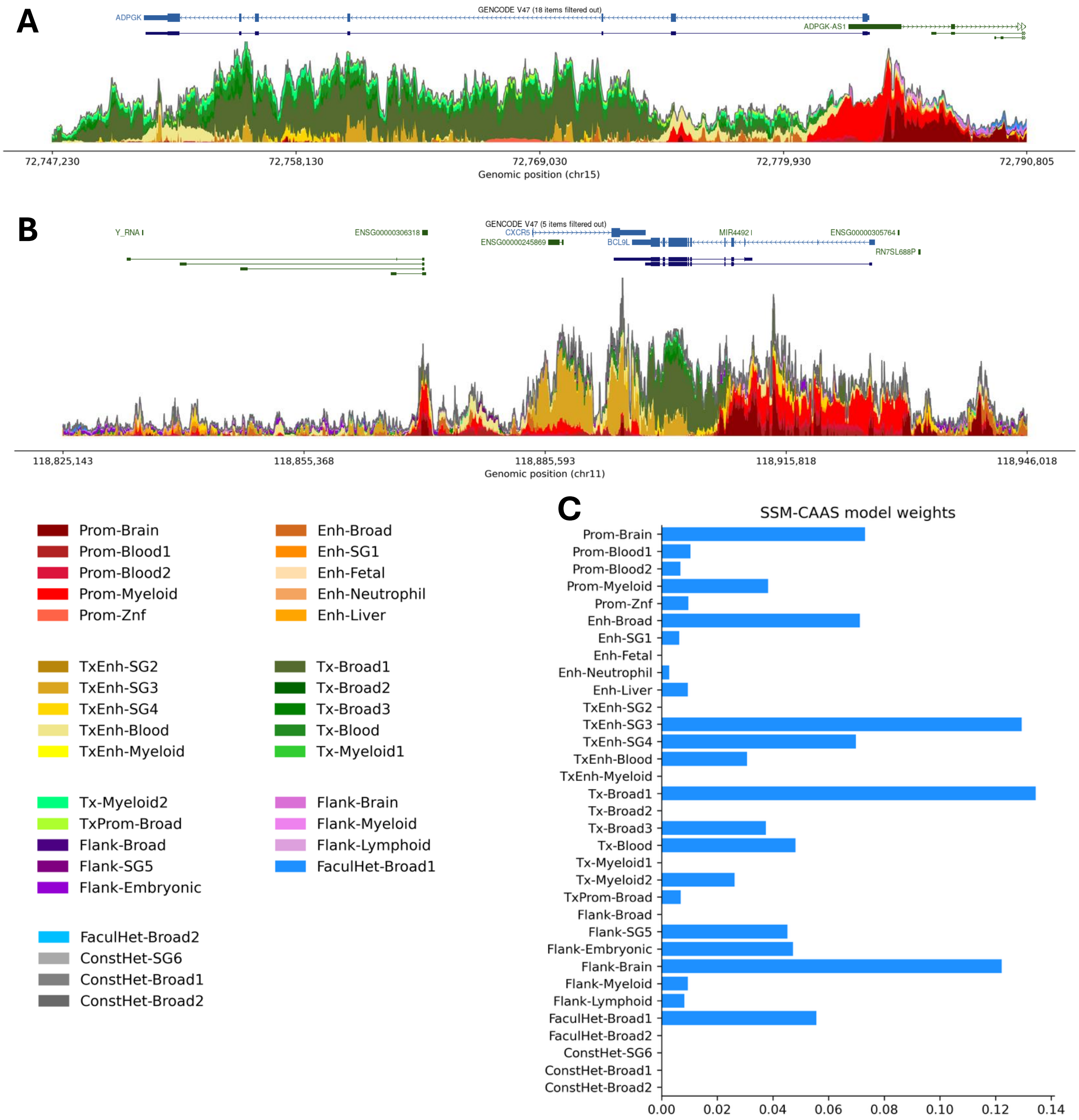


Figure S9: SSM-CAAS plot summarizes epigenomic signals at a given locus. **A** CAAS plot for the ADPGK locus. **B** CAAS plot for the CXCR5 and BCL9L loci. **C** The weights for the pan-cell type features in the SSM-CAAS regression model.

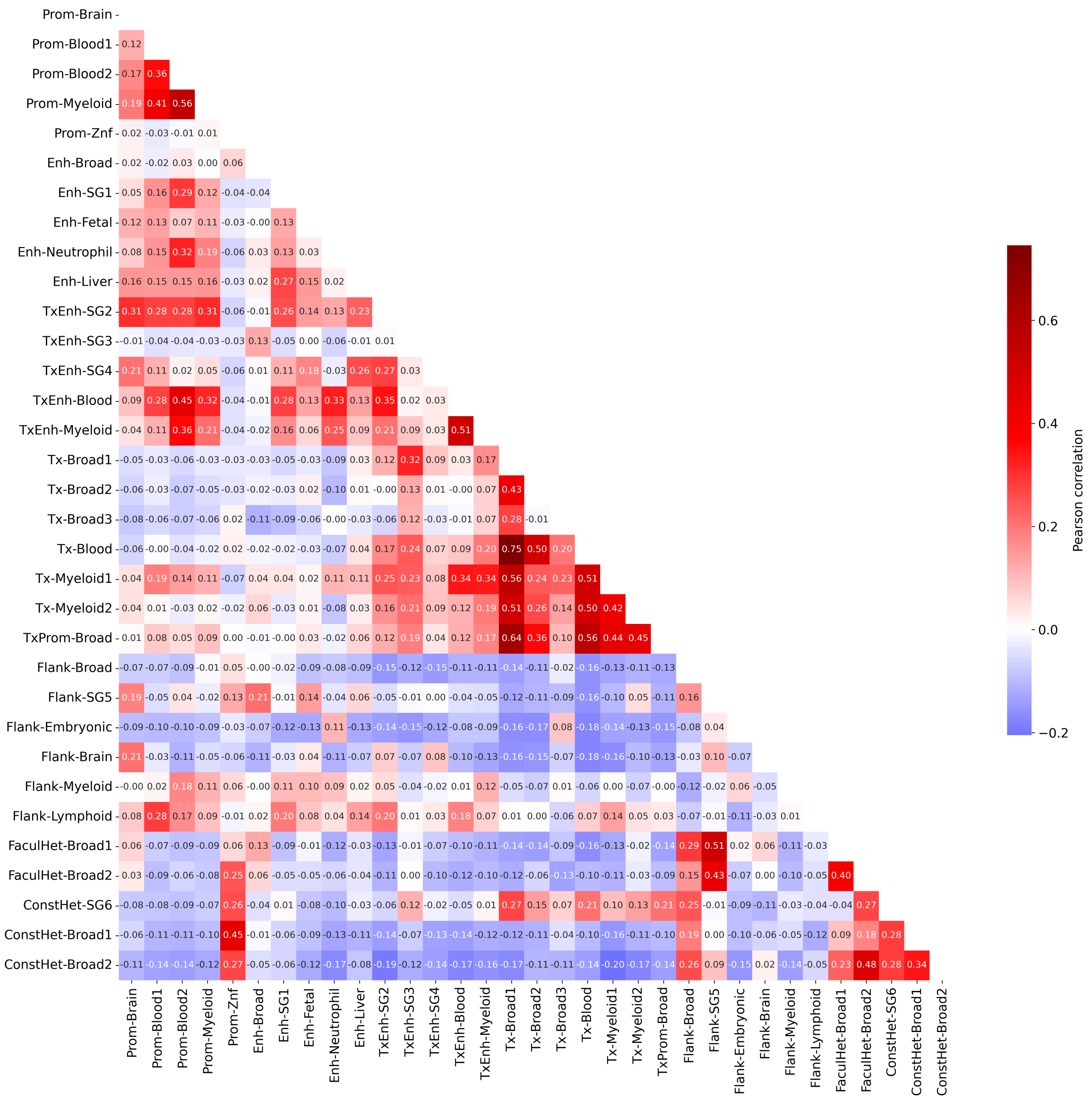


Figure S10: Correlation of pan-cell type features with each other.
